# Anterior cingulate cortex neurons in macaques encode social image identities

**DOI:** 10.1101/2023.11.10.566537

**Authors:** Joseph Simon, Erin L. Rich

## Abstract

The anterior cingulate cortex gyrus (ACCg) has been implicated in prosocial behaviors involving complex reasoning about social cues. While this indicates that the ACCg is involved in social behavior, it remains unclear whether ACCg neurons also encode social information during goal-directed actions without social consequences. To address this, we assessed how social information is processed by ACCg neurons in a reward localization task. Two rhesus monkeys performed the task using either social or nonsocial visual guides to locate rewarding targets. We found that monkeys can use both sets of guides, and many neurons in the ACCg distinguished social from nonsocial trials. Yet, this encoding was no more common in ACCg than in the prearcuate cortex (PAC), which has not been strongly linked to social behavior. However, unlike PAC, ACCg neurons were more likely to encode the unique identity of social visual guides compared to nonsocial, even though identity was irrelevant to the reward localization task. This suggests that ACCg neurons are uniquely sensitive to social information that differentiates individuals, which may underlie its role in complex social reasoning.

## Introduction

Processing information with respect to social context is a fundamental component of social cognition ^1–3^. In humans and nonhuman primates, this ability allows individuals to navigate social interactions and learn about their environment from others ^4,5^. To date, it remains unclear whether social information processing relies on unique computations or distinct neural circuits, or whether it employs general functions that also mediate complex behavior in nonsocial contexts ^6,7^. Given the prevalence of social information in daily life, it is important to understand the underlying neural codes governing its use.

In primates, the medial frontal cortex, particularly the anterior cingulate cortex gyrus (ACCg), has consistently been implicated in social information processing ^8–11^. Neuroimaging in humans has found activation in the ACCg when participants estimate the probability that a social partner is trustworthy ^12^, the similarity of others’ political beliefs or preferences to their own ^13^, or the volatility of information coming from a confederate ^4^. In monkeys, neurons in the ACCg encode the receipt of reward for both themselves and others ^10^, and lesions of this region reduce the tendency to form prosocial preferences ^9^. Together, these data indicate that ACCg plays an important role in producing socially appropriate behavior that takes others into account. However, compared to self-oriented goal-directed choices, these types of social decisions typically have added complexity, such as ambiguity about the states, beliefs, or actions of others. Overlapping regions of ACCg are also activated when making nonsocial decisions in complex environments, for instance when humans adapt their behavior based on uncertain evidence ^14^, or when monkeys make decisions that weigh costs versus benefits ^15^. Since the ACCg is implicated in both social reasoning and more general cognitive control processes, one possibility is that its involvement in social situations relates to the complexity of these tasks rather than the social nature of the task per se. Therefore, we aimed to study ACCg involvement in goal-directed behaviors of similar complexity that vary only in the involvement of social versus nonsocial information.

A natural social cue that can guide decisions is directed gaze ^2,16,17^. Recognizing that another individual is attending to an object or location in the environment can shift one’s attention toward the same direction, a behavior called gaze following. Gaze following can impact future decisions related to the information in the focus of shared attention ^18,19^, and is a conserved trait across primate species, being found in both adult and infant humans as well as monkeys ^17,20–22^. This behavior is also altered in some cases of neurodivergence. For example, the ability to follow a sender’s gaze can be affected in individuals with autism ^23^. Whether gaze following deficits represent unique alterations in social processing or are part of larger deficits in attentional, orienting, or salience detection mechanisms is unclear.

To better understand whether unique neural processes are engaged by social cues that inform decisions, we developed a reward localization task in which rhesus monkeys use gaze direction of a social cue or a nonsocial cue to select a visual target to get a reward. This task allowed us to determine whether ACCg neurons respond to social information in the absence of potential social outcomes (i.e., prosocial influences). In addition, our task varied whether the cues contained information about the location of the rewarded target. To determine whether encoding of social information is unique to the ACCg, we contrasted these neural responses to those in the prearcuate cortex (PAC), in and around the frontal eye fields. This region is not strongly implicated in social cognition, but rather in visual attention and planning upcoming eye movements ^24,25^, which are also important aspects of this task. Our results show that neurons in both regions differentiate social from nonsocial cues. However, the ACCg, but not PAC, showed a tendency to encode social identity information. This occurred despite social identity being irrelevant to solving the task. These results add to the growing body of literature implicating the ACCg in uniquely processing social information and suggest that identity encoding could underly its role in complex social reasoning.

## Results

### Task Performance

Previous research has shown that monkeys are able to follow the gaze of conspecifics ^17^. Here we tested whether monkeys could use an image of a conspecific directing their gaze to select a rewarded target. We used image sets of the same monkey face gazing to the left, right, or neither direction. Additionally, we assessed whether there were differences between this socially guided behavior and the same behavior guided by nonsocial cues in the form of arrows (**Figure 1**). Trials occurred in four blocks. One block contained all nonsocial image sets, and the others contained image sets of 2 monkey faces each. Each block included right, left, and nondirectional images counterbalanced and randomly selected. To begin each trial, monkeys had to hold a touch-sensitive bar and fixate a point on the task screen until an image appeared for 1000ms. Subjects could freely view the image if they continued to hold the bar. Then two identical green squares were shown on the right and left, and monkeys had to choose the correct target to receive a fruit juice reward (**Figure 2**). If the image did not indicate a direction, the squares were different colors (red and purple) and the monkey had to select the purple square to receive a reward. To make a choice, monkeys fixated the desired square and simultaneously released the bar. Data include 31,953 trials over 39 sessions for Monkey N, and 24,641 trials over 30 sessions for Monkey J.

**Figure 1:**
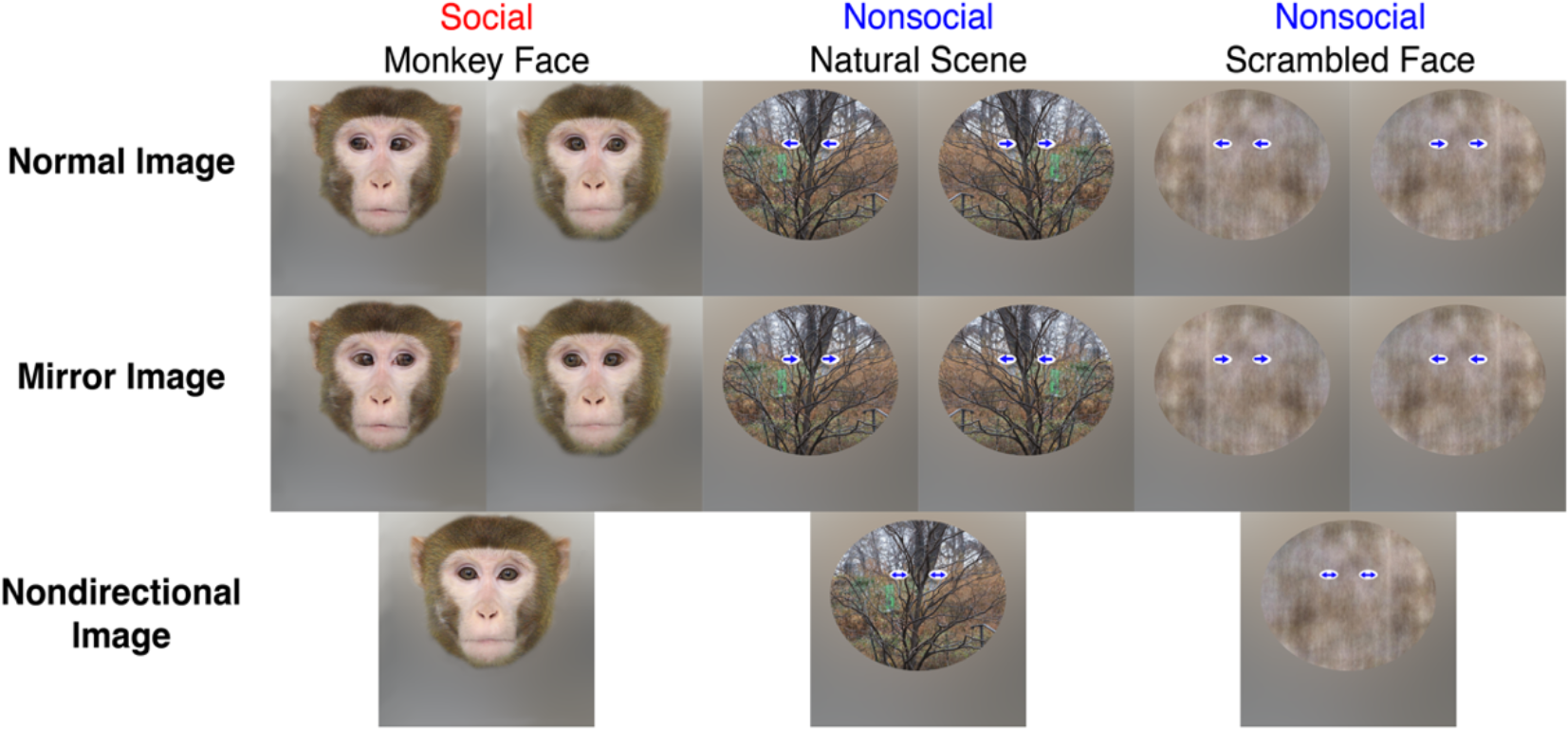
Visual guides. Example image sets for different types of visual guides. *Top row*: Visual guides with the same monkey gazing left or right (social), a natural scene with arrows pointing left or right (nonsocial), and a scrambled monkey image with arrows pointing left or right (nonsocial). *Middle row*: The same visual guides as the top row, but with eye gaze or arrow directions reversed. *Bottom row:* examples of visual guides of each type that do not give any directional information.

**Figure 2:**
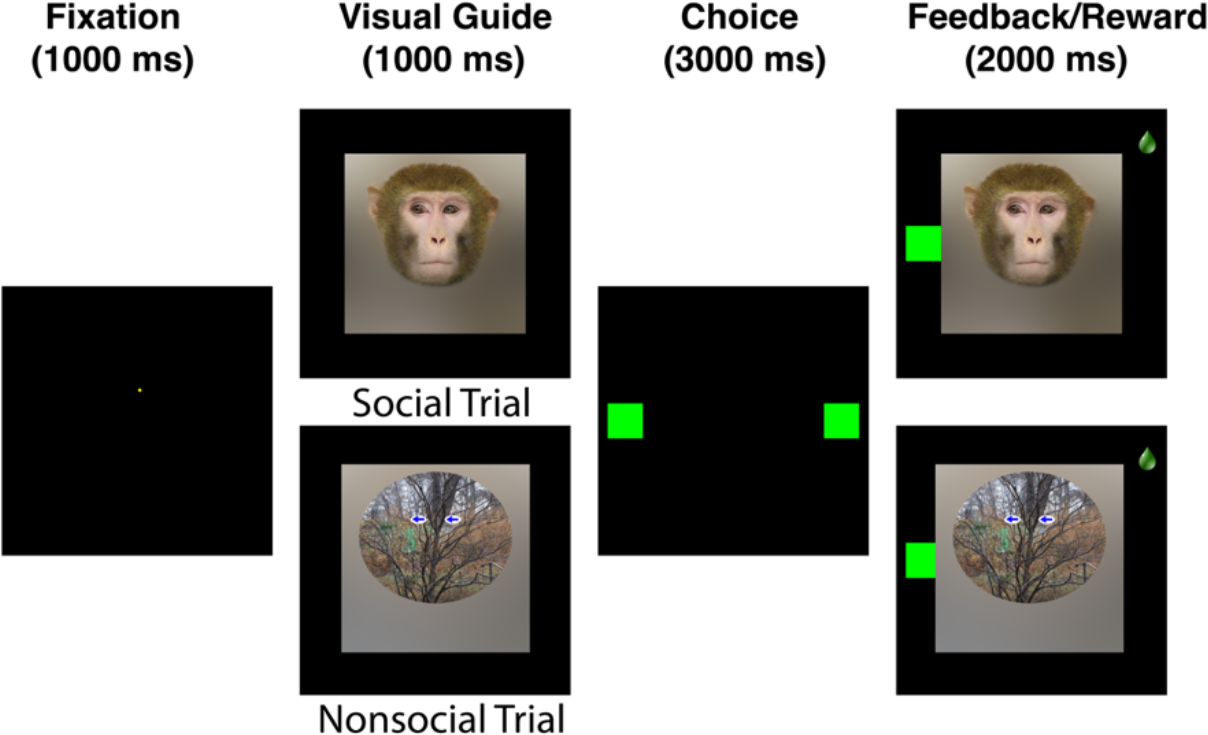
Behavioral task. Each trial required monkeys to fixate a point on the screen for 1000ms to begin.They were then shown either a social (*top*) or nonsocial (*bottom*) visual guide. Following visual guide presentation, two identical green squares appeared (or one purple and one reo square on nondirectional trials; not shown) Monkeys were given a juice reward if they choose the target indicated by the visual guide.

Given that the visual guides (i.e., eye gaze and arrow direction) conveyed the same information, we predicted that monkeys should be able to use either cue type to perform the task. We found that both monkeys performed significantly better than chance (> 50%) for both types of visual guides (**Figure 3**). Interestingly though, they both performed worse when using social guides, compared to nonsocial guides. A logistic regression analysis including context (social vs nonsocial), information source (directional visual guide vs target color), and gaze direction (left vs right), found a significant main effect of context in Monkey N (t_1,31953_ = -2.49, p *<* 0.05), and a trend toward significance in Monkey J (t_1,24641_ = -1.85, p = 0.065), with better performance on nonsocial trials in both cases. Given Monkey J’s high performance overall, the difference is likely a ceiling effect. Despite this asymmetry, however, performance was high in both contexts, demonstrating that monkeys can use gaze information from static social images to guide choice behavior.

**Figure 3:**
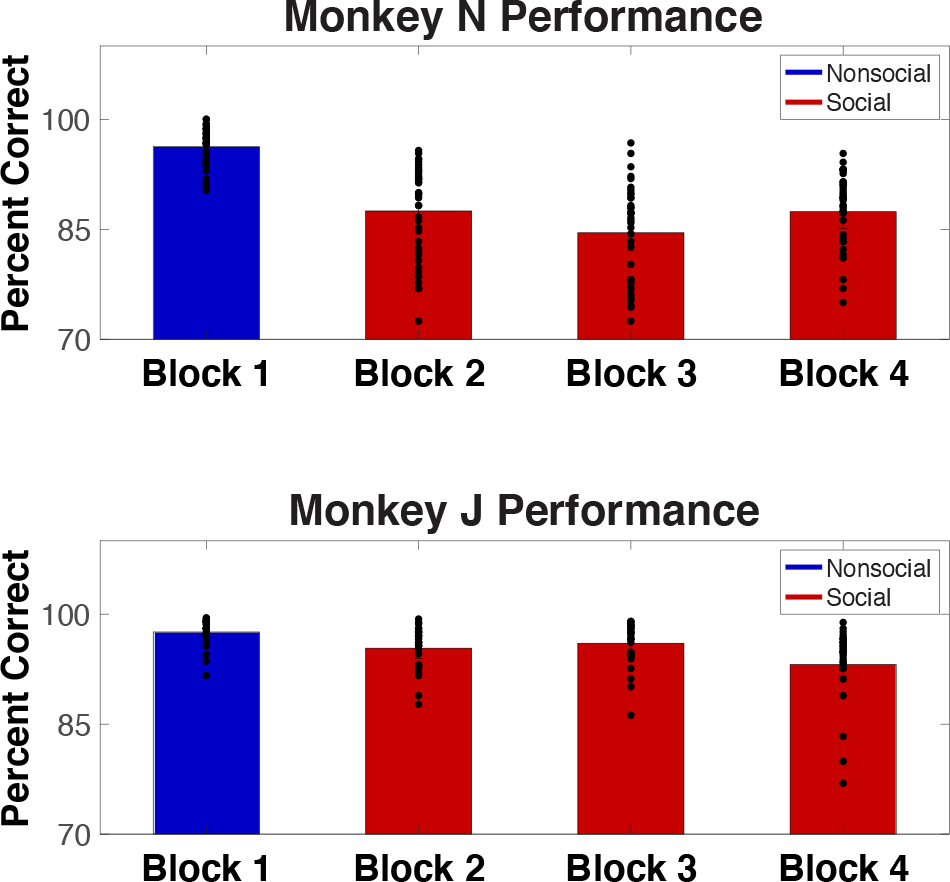
Behavioral performance across blocks. Percent of trials in which the correct target was selected for Monkey N and Monkey J. Block 1 (*blue*) includes all nonsocial trials and Blocks 2-4 (*red*) arc social trials. Dots indicate average performance per image set in each block.

### Gaze Behavior

Our task design allowed the monkeys to freely view the visual guides, so we quantified their natural viewing behavior to determine whether attending to a stimulus increased their ability to gather information and make better choices. In general, monkeys spent the most time looking at either the eyes or arrows of the visual guides, or at the cued target location (**Figure 4**). To determine whether performance was predicted by the amount of time monkeys looked at the discriminative region of the visual guides, we quantified the total time on each trial that their gaze fell within a *±* 5 visual degrees radius of the center of the eyes or arrows during correct and incorrect trials for both contexts. Multiple linear regressions determined whether gaze durations varied with performance (correct vs incorrect), context (social vs nonsocial), information source (directional guide vs target color), and the interaction of these parameters (**Figure 5**). We found a main effect of performance for both subjects (monkey N: t_31953_= 4.54, p *<* 0.001, monkey J: t_24641_= 4.81, p *<* 0.001), with longer gaze times on correct trials. In addition, there were effects of information source (monkey N: t_31953_= -2.76, p *<* 0.01, monkey J: t_24641_= -2.74, p *<* 0.01), with longer gaze times on nondirectional trials in both animals. For monkey N, there were also interaction effects of performance *×* context (t_31953_ = -3.49, p *<* 0.001), information source *×* context (t_31953_= -4.69, p *<* 0.001), and a three-way interaction (t_31953_= 3.46, p *<* 0.001) performance *×* information source *×* context, such that the longest gaze times were for correct responses to noninformative social images. In Monkey J, there was an interaction effect of performance *×* information source (t_1,24641_= 3.13, p *<* 0.005), where gaze duration was longer for correct responses on informative trials. However, we did not find consistent effects of social vs nonsocial context, suggesting that the type of visual guide had little impact on gaze duration. Together, these data show that there are idiosyncrasies in how monkeys attend to each stimulus, but both monkeys looked at the stimulus longer during non-directional trials and correct performance consistently accompanied longer viewing times.

**Figure 4:**
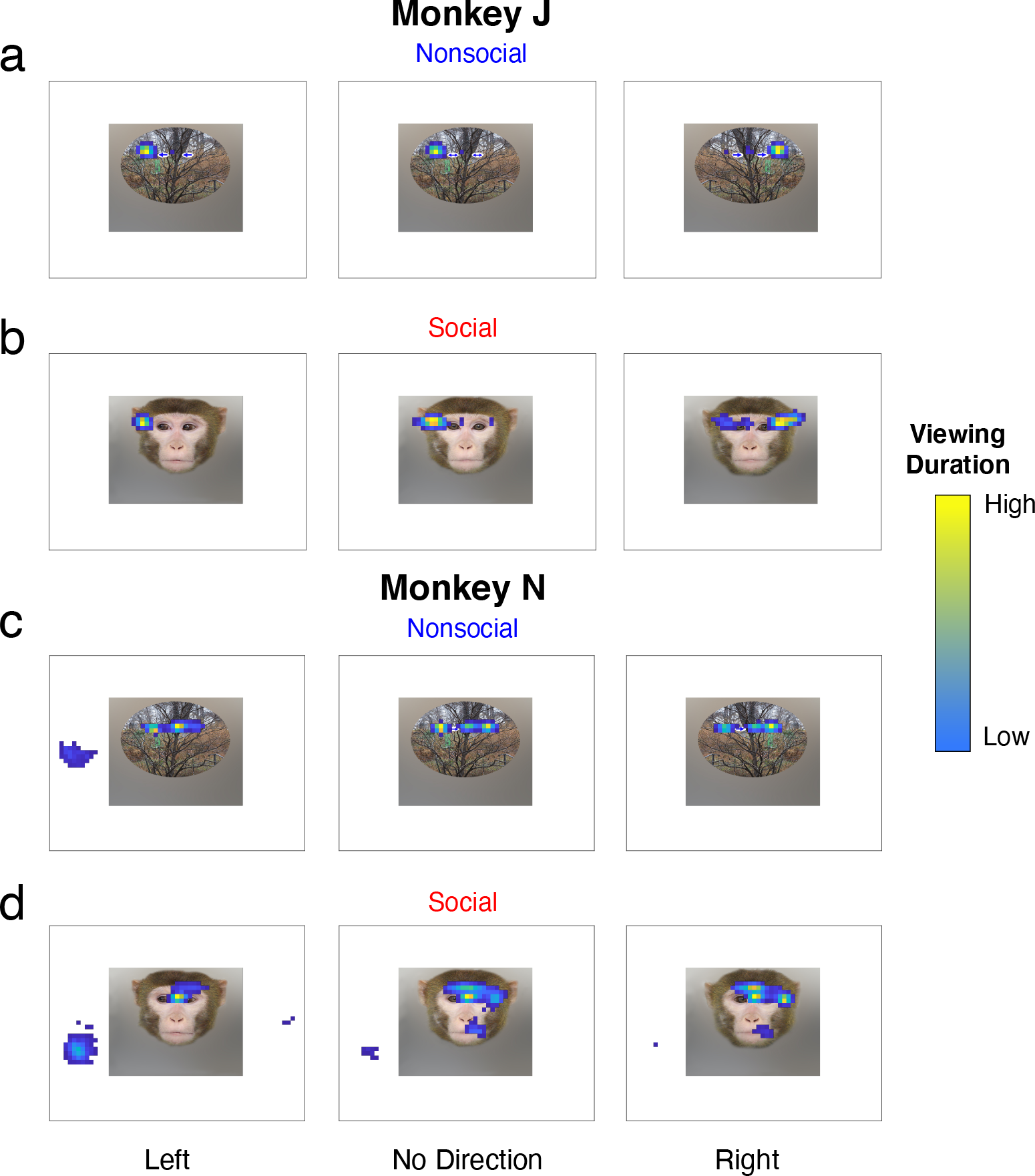
Examples of viewing behavior. Heatmaps display normalized viewing duration across all sessions for Left, No Direction, and Right visual guides, **(a, c)**, Nonsocial visual guide, **(b, d)**, Social visual guide.

**Figure 5:**
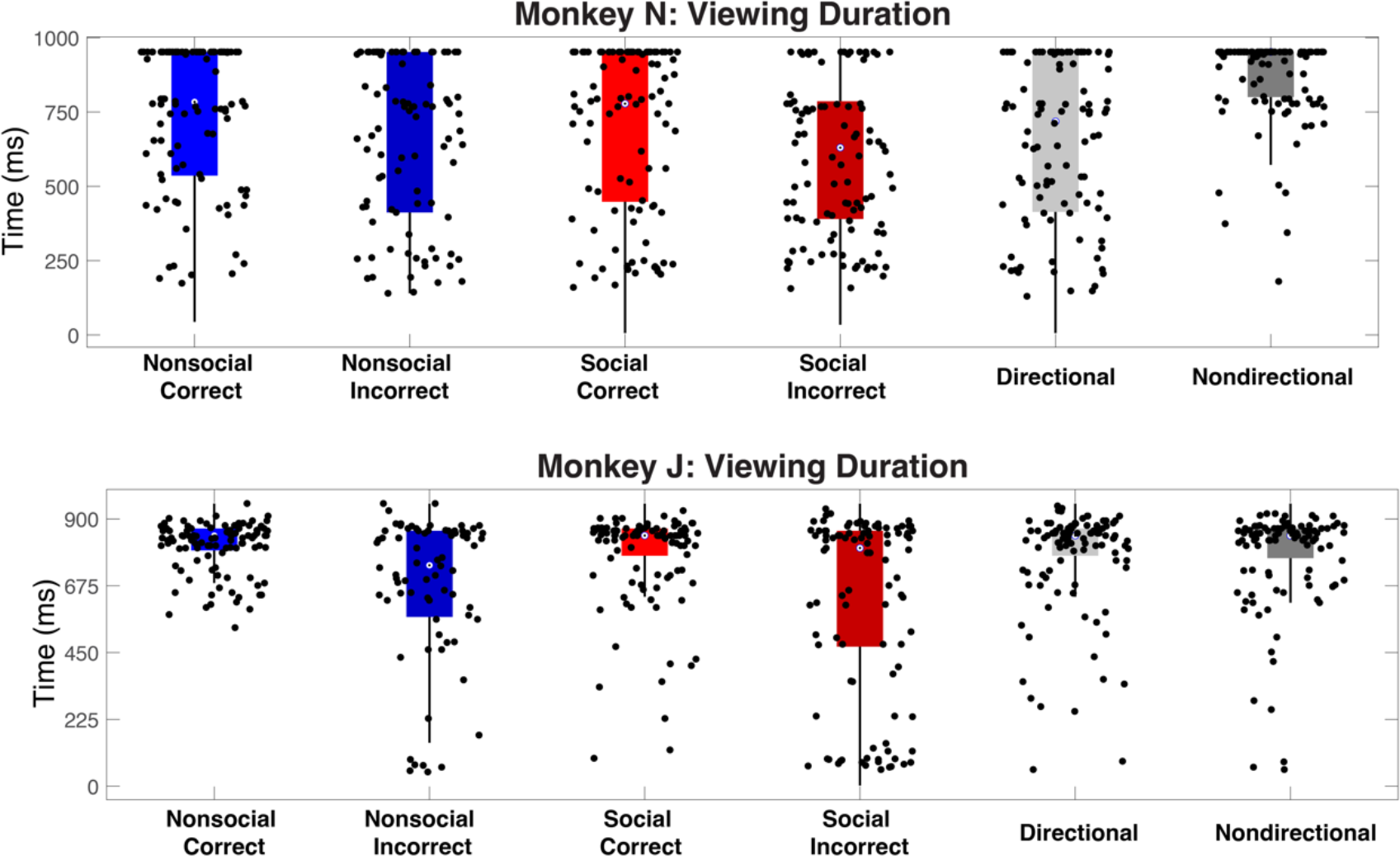
Gaze duration across task conditions. Viewing duration during visual guide presentation for correct and incorrect trials separated by social (*red*) and nonsocial (*blue*) context, also separated by information source (i.e., directional or nondirectional). Points show 100 representative trials, selected by down sampling the total trials in each condition. The maximum time was the duration of the visual guide presentation (1000ms).

### Neurophysiology

We recorded 101 and 114 neurons from ACCg and 102 and 126 from the PAC in subject N and J respectively (**Figure 6a-b**). We found that many ACCg neurons differentially responded to social and nonsocial contexts (**Figure 7**). To quantify neural encoding in this task, we fit the firing rate of each neuron in 200-ms sliding windows with a multiple regression model (Methods). Most neurons encoded context between the onset of the visual guide and 500-ms post onset in both ACCg and PAC (**Figure 7b**). A small number of neurons in the ACCg encoded context before image onset, which was possible because the cues appeared in blocks, so monkeys could have anticipated whether they would see a social or nonsocial image. To assess the tendency for neurons to respond more to social vs nonsocial cues, we calculated the average regression weight on the context predictor in the first 500-ms following stimulus onset for each neuron. Positive weights indicated greater firing rates for social stimuli, while negative indicated greater firing rates for nonsocial stimuli. Within the ACCg, similar proportions of neurons had positive and negative weights (50% Monkey N, 56% Monkey J, positive beta weights), and binomial tests confirmed that ACCg had no bias toward stronger responses for a particular context (p = 0.83). Within the PAC, there was also no overall bias toward stronger responses for a particular context (binomial test, p = 0.42). In Monkey J, there were similar proportions of neurons that responded to each context (52% for positive). Monkey N showed some difference in the relative proportion of selective neurons (13% positive), but binomial test showed that this was not statistically significant (p = 0.07).

**Figure 6:**
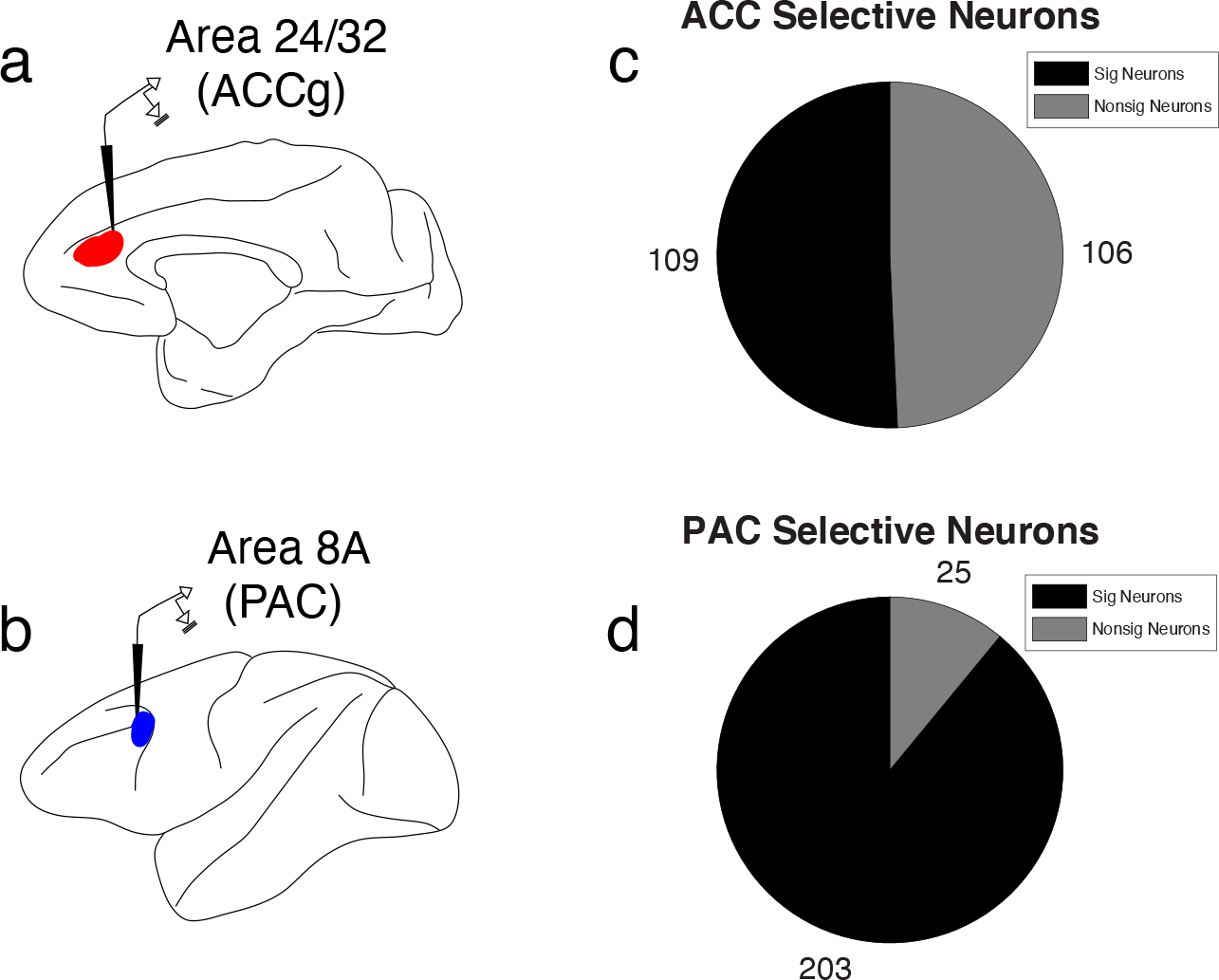
Neuron recording summary. **(a)** Target location in the ACCg. **(b)** Target location in the PAC. **(c-d)** Number neurons across both monkeys that were selective for one or more task variables (black) in the ACCg and PAC.

**Figure 7:**
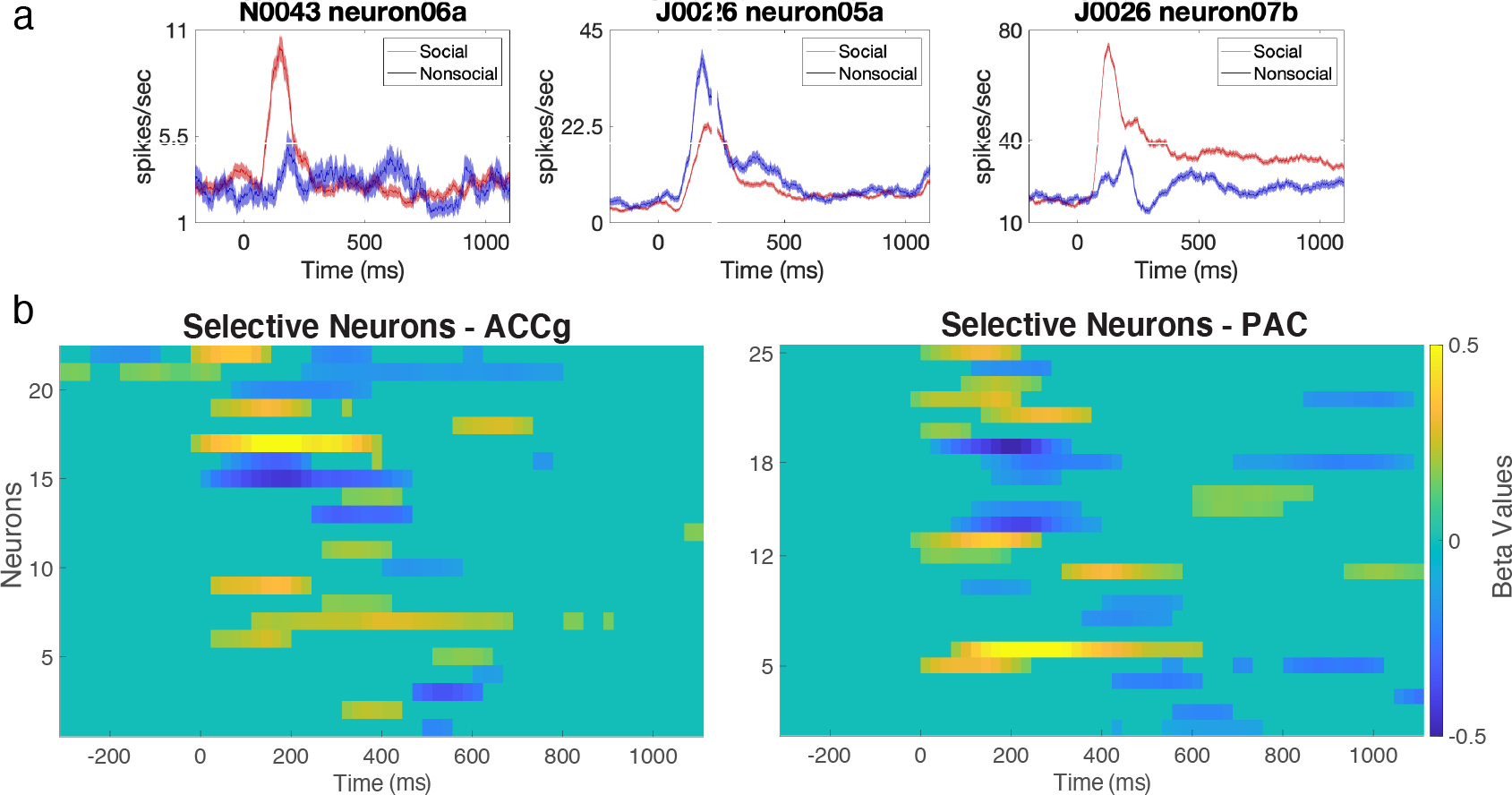
Neurons encoding social versus nonsocial contexts. **a)** Example responses of three ACCg neurons encoding social/nonsocial context during the visual guide period. **b)** regression weights for the context regressor for all neurons that demonstrated selectivity in a multiple regression during visual guide presentation in the ACCg (left) and PAC (right). Positive betas (warm colors) indicate stronger responses for social context, while negative betas (cool colors) represent stronger responses for nonsocial context.

Next, we assessed response latencies in both regions. We found that in PAC, but not ACCg, neurons that had higher firing rates for social contexts tended to have shorter latencies than those responding more to nonsocial contexts (Wilcoxon rank sum: PAC, z_pos-neg_ = -2.33, p = 0.021, ACCg, z_pos-neg_ = -1.02, p = 0.31). The reason for faster PAC responses to social cues is unclear, but it underscores that PAC neurons are differentially responsive to social and nonsocial images. Finally, we found no significant difference in the latency between regions (z_pos-neg_ = 1.19, p = 0.23).

Since there were not strong differences in how ACCg and PAC encoded social context, we assessed encoding of other variables in the task. To do this, we collapsed the sliding regression into four 1000-ms epochs, and found the number of neurons with significant encoding at any point in each epoch. During the fixation epoch, before the visual guide was presented, there was very little information encoded in either region, as expected (**Figure 8a**). When the visual guide appeared, the most common variable encoded by ACCg was context (10%), followed by information source (7%), followed by target location (3%) (**Figure 8b**). In addition, several ACCg neurons significantly encoded one or more unique social images (**Figure 8b**, red bars). Significance for the social identity regressors meant that a neuron responded to all image variations showing the same monkey in the visual guide. In the subsequent epochs surrounding choice and feedback, encoding of information source and target location increased (**Figure 8c-d**).

**Figure 8:**
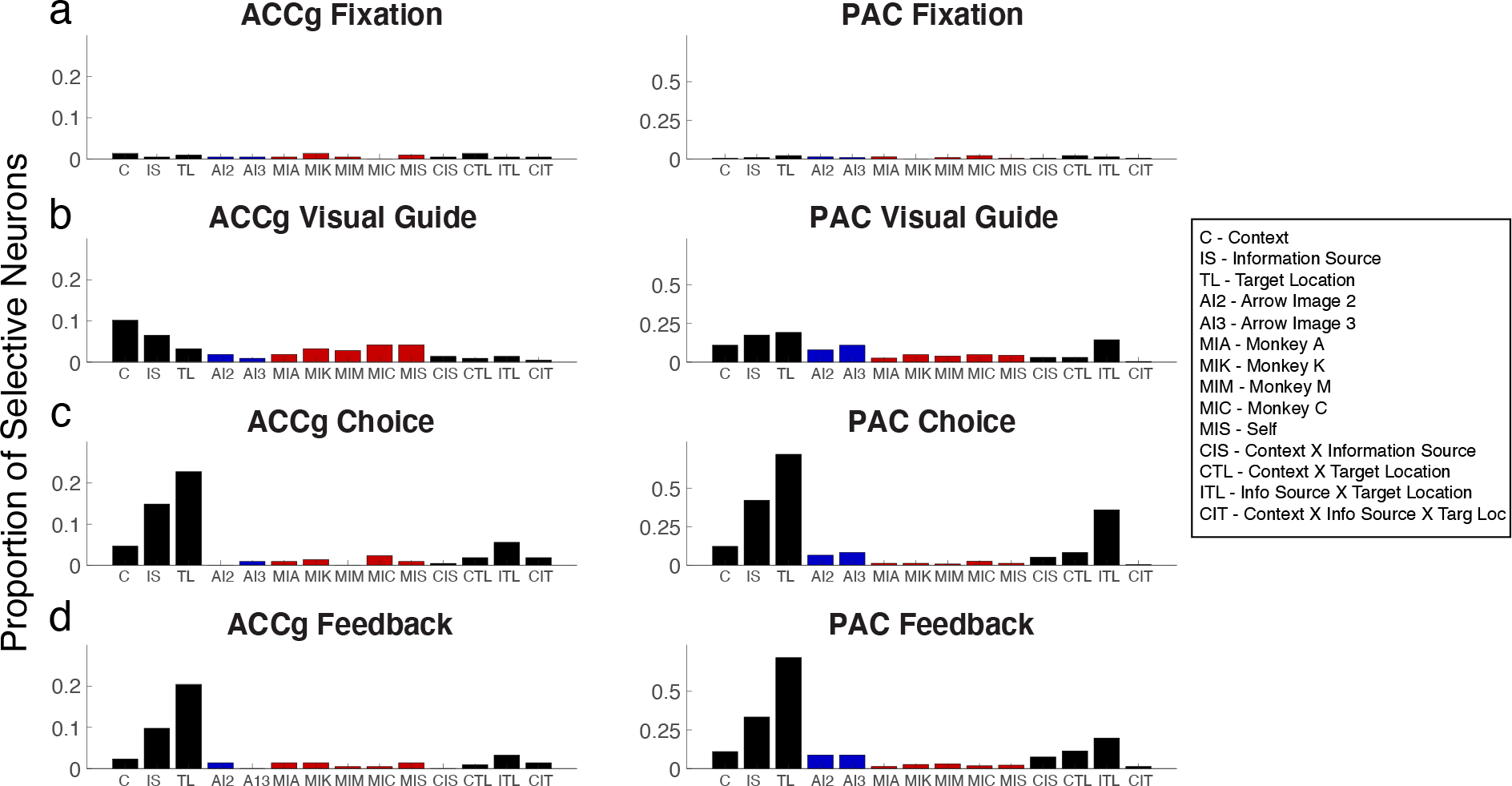
Single unit encoding in ACCg and PAC. Proportions of neurons in ACCg (*Left*) and PAC (*Right*) that significantly encoded task variables. Red bars indicate neurons significant for regressors that identified unique social identities. Blue bars show neurons significant for regressors that similarly identify each non-social image.

To determine whether these responses are unique to ACCg, we contrasted our results with neurons in the PAC, whose function is more related to visual attention and planning ^25^. Overall, larger proportions of PAC neurons encoded task-relevant information (**Figure 6**, right), but the ratios of encoded variables were comparable to ACCg, particularly in the choice and feedback epochs (**Figure 8 c,d**). Here, the largest proportion of neurons encoded gaze direction, followed by information source (directional image vs target color). In PAC there were also a large proportion of neurons that encoded the interaction of information source and target location (**Figure 8b-d**). The clearest distinction between regions was found during the visual guide epoch. In contrast to ACCg, the most prevalent encoding in PAC was target location (19%), consistent in PAC’s role in visuospatial attention, followed by information source (18%), followed by the interaction between target location and information source (14%) (**Figure 8b**). In addition, we observed more PAC neurons that responded to unique nonsocial images, compared to social images, the opposite pattern of that found in ACCg neurons.

To quantify the tendency of ACCg neurons to encode unique social identities and PAC to encode nonsocial images, we performed a *post-hoc* analysis. Binomial tests compared the proportion of neurons significantly encoding unique social images during the visual guide epoch to the proportion encoding unique nonsocial images, separately for both areas. To correct for the fact that there were more unique social images than nonsocial images, we normalized the neuron proportions by computing the average number of significant neurons per nonsocial image and multiplied this by the total number of social images. We found that a higher proportion of ACCg neurons encoded unique social images (binomial test, p ≤ 9.5×10^-6^, **Figure 9a**) and this was consistent in each monkeys’ data tested separately (Monkey N, p = 4.4×10^-5^, Monkey J, p = 0.012). We assessed the location of significant neurons to determine whether social identity neurons were anatomically clustered in the ACCg, however, we did not observe a definitive patten (**Figure 10**).

**Figure 9:**
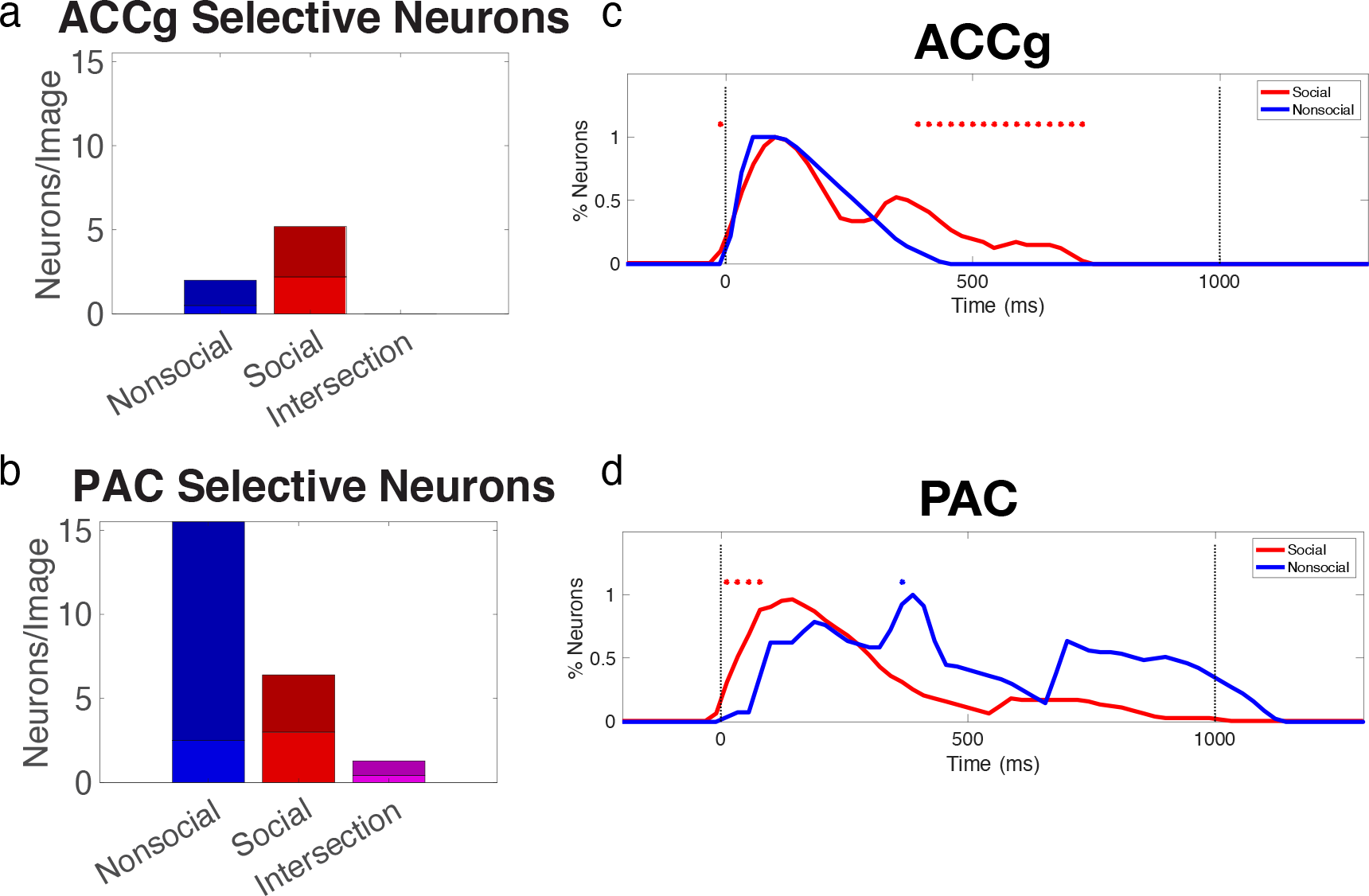
ACCg neurons encoding social identities, **(a-b)** Number of neurons per image encoding social identities only (red), nonsocial identities only (bkic), or both (purple) in the ACCg (a) and PAC (b). **(c-d)** Time course of responses relative to the appearance of the visual guide among significant neurons of each type in the ACCg (c) and PAC (d). Permutation tests determined significance. *p<0.05.

**Figure 10:**
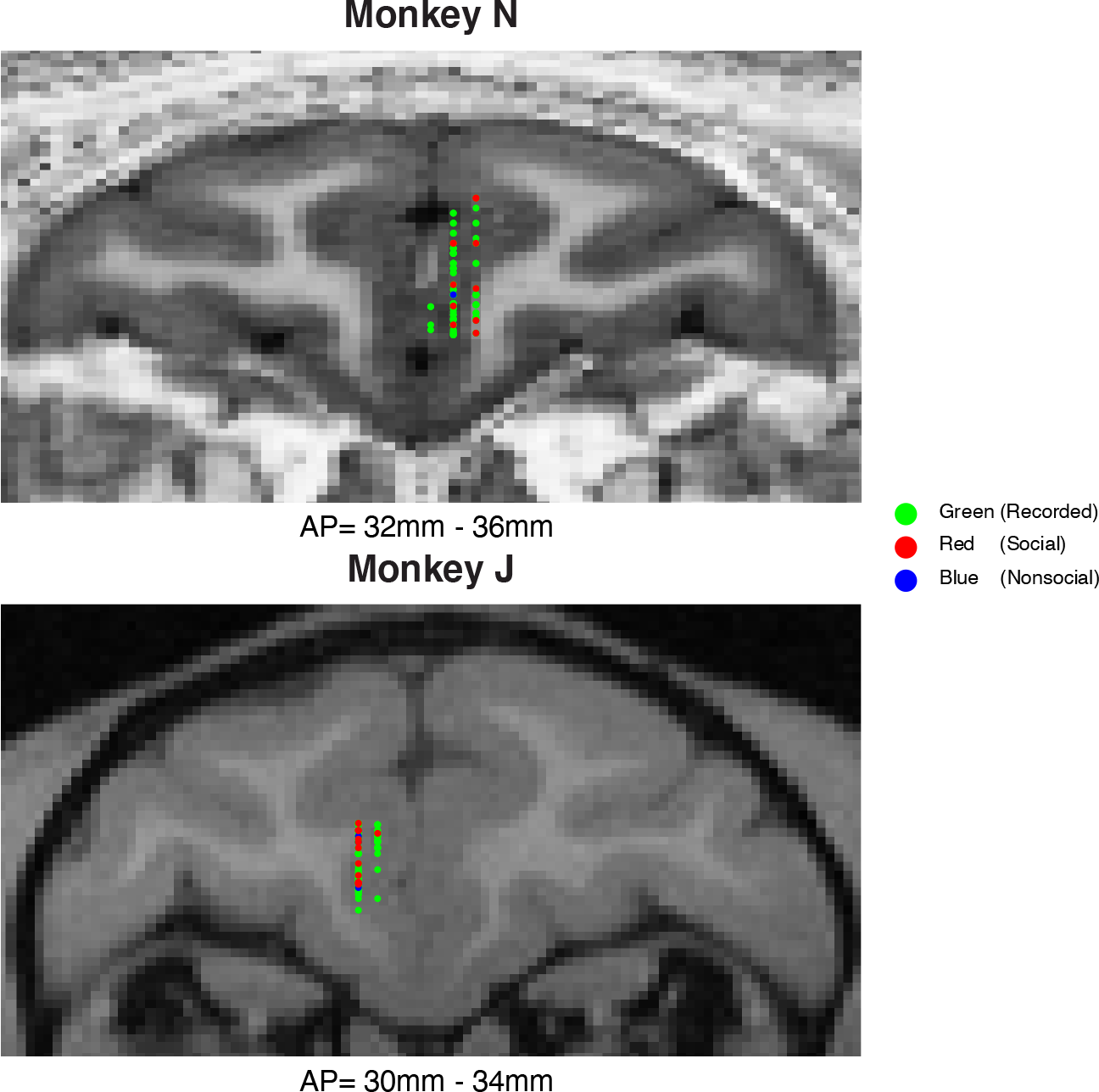
Neuron Locations. Coronal MR images though the Prefrontal Cortex. Images show the mid-point, in the anterior-posterior (AP) dimension, of the recording field in each subject, with all recorded neurons projected onto this slice. Note that this results in some recording locations appearing to fall slightly outside of gray matter boundaries. Colored dots represent recorded neurons: *Green*, all recorded neurons, *Red*, neurons that were significant for social image identity, *Blue*, neurons significant for nonsocial image identity.

We next assessed social identity neurons in the PAC. In contrast to ACCg, the encoding bias was in the opposite direction. A higher proportion of PAC neurons significantly encoded unique nonsocial images compared to social (**Figure 9b**), although this finding was driven by Monkey J (Monkey N, binomial test p = 0.45, Monkey J, p = 1.7×10^-14^). Since more than one of the identity regressors might significantly explain neuron firing (i.e., a neuron may respond to more than one image set), we quantified the number of neurons that reached significance for both a social and a nonsocial image (‘intersection’). We found that only PAC had neurons that responded to both a social and a nonsocial image, whereas in ACCg neurons mainly encoded social images only, and single neurons never encoded a social and a nonsocial image (**Figure 9a-b**). Finally, we assessed the time course of encoding each image type. Identity selective neurons responded with similar time courses in ACCg, regardless of the type of image they encoded, but social identity neurons continued to be active significantly longer during stimulus presentation (**Figure 9c**). On the other hand, PAC neurons encoded social images slightly earlier than nonsocial images (**Figure 9d**). This is consistent with our earlier finding that neurons in PAC that responded to social over nonsocial contexts tended to respond earlier. Together these data could suggest that social information reaches the PAC faster than information from other sources, suggesting a distinct circuit.

## Discussion

Social interactions are important to our daily lives, and the ability to track an agent’s identity is a fundamental component of normal social interactions. Here, we found that a small but significant proportion of neurons in the ACCg encoded the identity of conspecifics in social images, but not identities of nonsocial images. This occurred in a task where social or nonsocial images played identical roles in guiding behavior, and where the identity of the monkeys had no relevance to accurate performance. Similar selectivity was not found in a frontal region more involved in visuospatial attention, the PAC. This suggests that the tendency for single neurons to respond mainly to images of a particular monkey, rather than a familiar image in general, is a unique property of the ACCg that may be important in more complex social tasks. In particular, it suggests that ACCg may be involved in parsing and/or tracking the identities of social partners.

Our results are in line with other recent evidence showing that ACC processes social identities ^18,26^. For instance, in monkeys engaging in a 3-way prisoners dilemma task where they had to choose which partner to give a reward to, the animals tracked the decisions of each partner and used this information to decide how to interact with them ^18^. Neurons on the dorsal bank of the ACC sulcus encoded information about partner identities as well as their past behaviors, all information that could be used to make socially-relevant choices. Our study provides an important compliment to these results, showing that neurons that encode social identities are not sensitive to the identity of nonsocial images when making identical decisions. Together, these studies support the notion that the social identity coding we observe in ACC is directly linked to social decision making.

In humans, multivoxel activity patterns in a similar region of ACC also differentiate the unique identities of human photos ^26^. Similar to our study, social images were used to convey task information, but unique identities were associated with different degrees of accuracy of information they provided. In that case, the same region of ACC tracked confidence in the advisor, again suggesting that identity encoding is important for guiding future behavior. In our task, there was no difference in the predictive ability of different social images, nor nonsocial, and in fact monkey identity was entirely incidental to the task. Despite this, neurons in the ACCg still differentiated image identity when it came from a social source. This suggests that responding to social identities may be an automatic function of the ACCg, that occurs even when it is not relevant to ongoing behavior.

In broader studies of social cognition, the ACCg in both monkeys and humans has been found to play an important role in social decision-making. For example, the trust game ^27,28^ and ultimatum game ^29^ require participants to track individual identities or other schema that can be used as a basis for making judgements when interacting with their partner(s). Additionally, monkeys with circumscribed lesions to the ACCg cannot learn from vicarious reward delivered to a conspecific ^9^. And similarly, monkeys that watched partners perform a task were able to perform the same task significantly better compared to not observing a partner, and this ability was also linked to activity in the ACC ^30,31^. Therefore, there is strong evidence that ACC, and ACCg in particular, is important for social decision-making.

In contrast to ACCg, we found that some neurons in PAC encoded unique social images, but these were intermixed with similar or greater proportions (in the case of monkey J) of neurons encoding the identity of nonsocial images. Some PAC neurons even responded to more than one image type. This suggests that PAC neurons parse unique features of similar trials, but don’t preferentially encode social identities like ACCg neurons. Similarly, we found that many PAC neurons also distinguished social versus nonsocial contexts. This is in line with previous studies showing that these neurons can encode more abstract, contextual information ^25,32^, in addition to more concrete spatial or attentional information. Although we made efforts to match the size and location of the arrows to the eyes, contextual differences could still be driven by features of the task, such as the need to recall the rules of how to interpret eyes or arrows.

An important limitation of our study is that we used only female monkeys, and it’s not clear whether males would respond to social images in a similar manner. Previous studies have shown that dominance as well as the sex of the monkey can play a role in social behavior and decision-making ^33^. We suspect that these effects would be limited in our task, since it was designed to remove any social interaction or social decision-making, and instead focus on target selection for self-reward. Nonetheless, future studies will be needed to understand if there are any link to sex or dominance in ACCg activity related to social identities.

Overall, there is a growing body of literature implicating the ACCg, along with other regions, in a putative circuit for social decision-making ^6,34–36^. Our study expands on this by showing that the ACCg is uniquely sensitive to the identity of social images when using eye gaze to infer target information. An important component of social interactions is the ability to interpret gaze, including in relation to the actor being observed, and this aspect of our study could have implications for understanding impaired gaze following and social functioning in neurodivergent individuals ^37^. Of note, there are direct unidirectional connections between the ACCg and PAC, particularly the frontal eye fields ^38,39^, which may provide a circuit by which social information directs one’s own gaze. Beyond gaze, identity coding is important for myriad social behaviors, including pair bonding and group-level dynamics, and understanding how ACCg neurons interact with wider social-brain circuits will help reveal mechanisms that produce complex socially-appropriate behavior.

## Methods

### Subjects

We trained two female rhesus macaques *(Macaca mulatta)*, aged 6 and 9 years and weighing approximately 5.7 and 6.8 kg at the time of recording (Monkey J and N, respectively). Monkeys were socially housed with at least one companion monkey in their home cage with visual access to other monkeys in the colony. Each subject underwent cranial surgery to implant a head positioner for accurate eye-tracking and to minimize movement during neural recording, as well as a unilateral recording chamber positioned over either the left (Monkey J) or right (Monkey N) hemisphere. Monkey N had an acrylic headpost while Monkey J had a titanium headpost. Chamber dimensions were 35.19 mm × 27.14 mm, and placement was calculated using images obtained from a structural MRI scan at 3T for each subject. The chambers were positioned to allow access to both ACCg (areas 24/32) and the PAC (area 8A). All procedures were in accordance with the National Institute of Health guidelines and recommendations from the Icahn School of Medicine at Mount Sinai Animal Care and Use Committee.

### Behavioral Task

Subjects sat in a primate chair and viewed a computer monitor. A touch-sensitive bar was affixed to the front of the chair within the monkeys’ reach. Eye movements were tracked with an infrared tracking system (ISCAN). Behavioral interfaces were controlled with NIMH MonkeyLogic software ^40^. To begin each trial, monkeys held the bar and fixed their gaze within 2° visual angle of a fixation cue for 1000-ms. The fixation cue was located at 5° above the origin to place the eyes in the center of the discriminative cues that were subsequently presented. Visual guides were approximately 10° *×* 10° in size. Following visual guide presentation, two identical green squares, 2° *×* 2° in size, were shown to the right and left. To discourage any reflexive actions (e.g., the monkeys responded as soon as the visual cue turned off) the start of the choice epoch was offset by steps of 250-ms between 0-1 second (i.e., 0-ms, 250-ms, 500-ms, 750-ms, 1000-ms). Once they made a choice, the unchosen square disappeared, and the visual guide reappeared along with their selected choice. Correct choices resulted in fruit juice reward; incorrect selections resulted in a 6-second timeout.

Two types of visual guides defined two different contexts, social and nonsocial. Social images were monkey faces and nonsocial images were arrows on a complex background. There were 9 image sets in total, consisting of 3 nonsocial sets, and 6 social sets (3 male monkey faces and 3 female monkey faces). Each set included otherwise identical images that indicated left, right, or neither direction. Each social image was a monkey face with neutral expression and gaze directed to the left or right, or forward-facing gaze (see Figure 1). The nonsocial images included pairs of arrows pointing to the left or right, or bidirectional arrows, set in white circles of approximately the same size and location as the monkey eyes, placed on a complex background. Backgrounds were either a natural scene, (i.e., trees in a park), or scrambled pixels of the monkey faces. The former was included because they are recognizable, natural images whereas the latter match the monkey faces in low level visual statistics. For all directional images (both social and nonsocial), images were digitally manipulated to flip, horizontally, the eye gaze or arrow background, while holding everything else constant. This was done to make sure that the monkeys only used gaze direction or arrow direction to select the target, and no other defining landmarks in the images. For nondirectional trials (i.e., eyes looking forward or double-headed arrows), the images did not indicate the correct target, so the targets consisted of a red and purple square instead of identical green squares, and the purple square was always correct. These trials were included to assess whether neurons encoded informative and non-informative visual guides similarly. Trials occurred in blocks. Monkeys finished a block by completing 25 trials correctly, at which point a new block was pseudorandomly selected, with the constraint that blocks never repeated sequentially. Sessions were completed after approximately 800 correct trials.

### Behavioral Analysis

Behavioral analyses were carried out separately for each subject. To quantify task performance, we used a logistic regression to predict correct or incorrect target selection from context (i.e., social versus nonsocial), information source (i.e., directional guide versus target color), cued direction (i.e., left versus right), and the interactions between these predictors.

Since monkeys free-viewed the visual guides, we also assessed how task variables influenced the amount of time they viewed the discriminative parts of the cues on each trial. To do this, we quantified the amount of time that the monkeys’ eyes fell within a radius of 5° of the center of the eyes or arrows in the visual guide, and used a multiple regression analysis with the parameters above, with the addition of performance (i.e., correct trials versus incorrect trials), to predict viewing time.

### Neurophysiological Recording

We used standard methods for acute neurophysiological recording, which have been described in detail elsewhere (see Lara et al., 2009). Briefly, in each recording session, we simultaneously recorded from 3-12 tungsten microelectrodes (FHC, Inc) or 16-channel linear arrays (V-probes, Plexon). A guide tube was placed to penetrate dura for each electrode. Since Monkey N had an acrylic headpost, we placed an empty guide tube positioned outside the recording field for ground. Monkey J had a titanium headpost, which was used for grounding. Each recording day, electrodes were manually lowered with custom-built microdrives to a target depth. We calculated target depths stereotaxically using 3T structural MRI images. Once lowered to desired depth, fine adjustments were made to isolate waveforms from single neurons. We recorded from all well-isolated neurons in target areas, resulting in a random sampling of neurons. Waveforms were acquired and digitized using an acquisition system (Ripple Neuro) and saved for off-line analysis. Single units were isolated offline with manual cluster cutting, performed using Offline Sorter (Plexon).

### Neurophysiological Analysis

To capture dynamics of encoding across a trial, we fit a multiple regression model (Eq. 1) to the firing rates of each neuron in sliding 200-ms windows. Model predictors were context (social versus nonsocial), information source (directional guide versus target color), cued direction (left versus right), the interactions between all predictors, and a matrix of image identities coded as one-hot vectors, using one social and one nonsocial image as an uncoded reference.

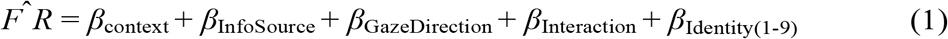

We analyzed neural activity during four trial epochs: fixation, visual guide on, choice, and feedback. All analysis epochs were chosen *a priori*, based on task events. The first window in each epoch started 200-ms before the onset of the relevant event, then analysis windows were stepped forward by 20-ms. This process was repeated through 1200-ms after epoch onset. To establish a significance criterion, we ran this same sliding regression over the fixation epoch, when nothing was present on screen and selected a criterion that resulted in ≤ 5% false discovery rate for information source and cued direction, which could not be anticipated at this time. With this criterion, significant encoding was defined as p < 0.0005 for Monkey J and p < 0.003 for Monkey N, each for 4 consecutive time windows.

We quantified the observed difference between neurons selective for social and nonsocial identity using a binomial test (Eq. 2). The number of selective neurons per nonsocial image was multiplied by the total number of social images to normalize our groups. The binomial test was then used to compare the probability of observing the proportion selective social neurons to nonsocial neurons.

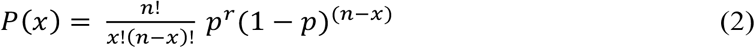

To capture differences in encoding probability between social and nonsocial visual guides, we performed a permutation test. We randomized neurons that encoded either social or nonsocial information using 10,000 permutations. We selected time bins that were greater than 95% of permutations.

## Acknowledgements

We thank Peter Rudebeck and Mark Baxter for comments on the manuscript, and Feng-Kuei Chiang with surgical assistance.

## Funding

NS125826 to JS.

## Notes

### Competing Interest Statement

The authors have declared no competing interest.

